# A Graph Extension of the Positional Burrows-Wheeler Transform and its Applications

**DOI:** 10.1101/051409

**Authors:** Adam M. Novak, Erik Garrison, Benedict Paten

**Affiliations:** Genomics Institute, University of California Santa Cruz, Santa Cruz, CA 95064, USA; Wellcome Trust Sanger Institute, Cambridge, UK

## Abstract

We present a generalization of the Positional Burrows-Wheeler Transform (PBWT) to genome graphs, which we call the gPBWT. A genome graph is a collapsed representation of a set of genomes described as a graph. In a genome graph, a haplotype corresponds to a restricted form of walk. The gPBWT is a compressible representation of a set of these graph-encoded haplotypes that allows for efficient subhaplotype match queries. We give efficient algorithms for gPBWT construction and query operations. We describe our implementation, showing the compression and search of 1000 Genomes data. As a demonstration, we use the gPBWT to quickly count the number of haplotypes consistent with random walks in a genome graph, and with the paths taken by mapped reads; results suggest that haplotype consistency information can be practically incorporated into graph-based read mappers.

## 2 Introduction

The PBWT is a compressable data structure for storing haplotypes that provides an efficient search operation for subhaplotype matches [2]. Implementations, such as BGT (https://github.com/lh3/bgt), can be used to compactly store and query thousands of samples. The PBWT can also allow existing haplotype-based algorithms to work on much larger collections of haplotypes than would otherwise be practical [4]. In the PBWT, each site (corresponding to a genetic variant) is a binary feature and the sites are totally ordered. The input haplotypes to the PBWT are binary strings, with each element in the string indicating the state of a site. In the generalization we present, each input haplotype is a walk in a general bidirected graph. This allows haplotypes to be partial (they can start and end at arbitrary nodes) and to traverse arbitrary structural variation. It does not require the sites (nodes in the graph) to have a biologically relevant ordering in order to provide compression. However, despite these generalizations, the core data structures are similar, the compression still exploits genetic linkage and the haplotype matching algorithm is essentially the same.

## 3 Definitions

We define *G* = (*V, E*) as a **genome graph** in a bidirected formulation [5,6]. Each node in *V* has a DNA-sequence label; a left, or 5′, **side;** and a right, or 3′, side. Each edge in *E* is a pairset of sides. The graph is not a multigraph: only one edge may connect a given pair of sides and thus only one **self-loop** can be present on any given side.

We consider all the sides in the graph to be (arbitrarily) ordered relative to one another. We also define the idea of the **opposite** of a side *s*, with the notation 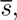 meaning the side of s’s node which is not *s* (i.e. the left side of the node if *s* is the right side, and the right side of the node if *s* is the left side). Finally, we use the notation *n*(*s*) to denote the node to which a side *s* belongs.

Within the graph *G*, we define the concept of a **thread**, which can be used to represent a haplotype or haplotype fragment. A thread *t* on *G* is a reversible nonempty sequence of sides, such that for 0 ≤ *i* < *N* sides *t*_2*i*_ and *t*_2*i*+1_ are opposites of each other, and such that *G* contains an edge connecting every pair of sides *t*_2*i*+1_ and *t*_2*i*+1_. In other words, a thread is a walk through the sides of the graph that alternates traversing nodes and traversing edges and which starts and ends with nodes. Note that a thread is reversible: exactly reversing the sequence of sides making up a thread produces an equivalent thread. We call a thread traversed in a certain direction an **orientation**.

We consider *G* to have associated with it a collection of **embedded** threads, denoted as *T*. We propose an efficient storage and query mechanism for *T* given *G*.

We call an instance of side in a thread a **visit**; a thread may visit a given side multiple times. Consider all visits of threads in *T* to a side *s* where the thread arrives at *s* either by traversing an edge incident to *s* (and not by traversing *n*(*s*)) or by beginning at *s*. For each such visit, take the sequence of sides coming before this arrival at *s* in the thread and reverse it, and then sort the visits lexicographically by these sequences of sides, breaking ties by an arbitrary global ordering of the threads. Then, for each visit, look two steps ahead in its thread (past *s* and 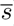), and note what side comes next (or the null side if the thread ends). After repeating for all the sorted visits to *s*, take all the noted sides in order and produce the array *B*_*s*_[] for side *s*. An example *B*[] array and its interpretation are shown in Figure 1. (Note that, throughout, arrays are indexed from 0 and can produce their lengths trivially upon demand.)

**Fig. 1.**
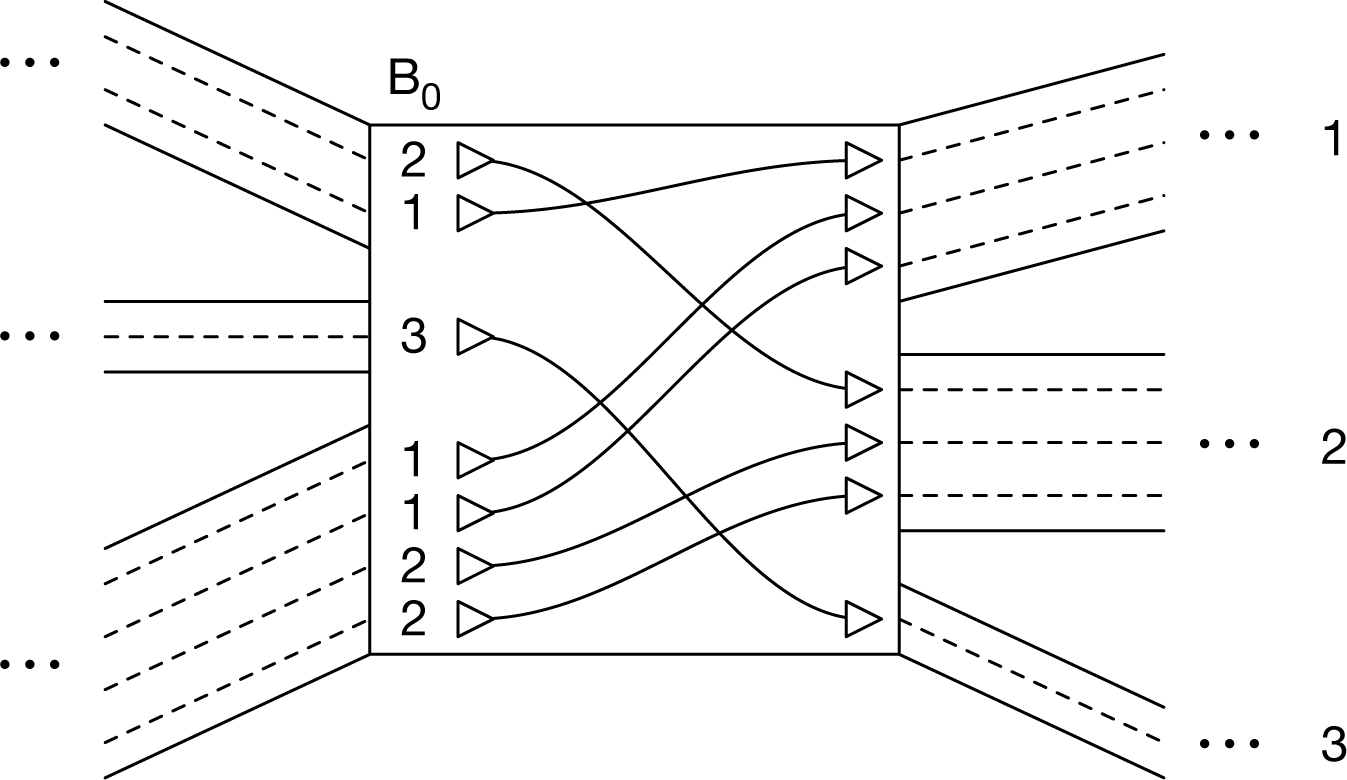
An illustration of the *B*_0_[] array for a single side numbered 0. Threads visiting this side may enter their next nodes on sides 1, 2, or 3. The *B*_0_[] array records, for each visit of a thread to side 0, the side on which it enters its next node. This determines through which of the available edges it should leave the current node. Because threads tend to be similar to each other, they are likely to run in “ribbons” of multiple threads that both enter and leave together. These ribbons cause the *B*_*s*_ [] arrays to contain runs of identical values, which may be compressed.

Each unoriented edge {s,*s’*} in *E* has two orientations (*s, s’*) and (*s’, s*). Let c() be a function of these oriented edges, such that for an oriented edge (*s’, s*), *c(s, s')* is the smallest index in *B*_*s’*_ [] of a visit of *s’* that arrives at *s’* by traversing {*s,s’*}. Note that, because of the global ordering of sides and the sorting rules defined for *B*_s’_[] above, *c*(*s*_0_, *s’*) will be less than or equal to *c*(*s*_1_, *s’*) for *s*_0_ <*s*_1_ both adjacent to *s’*.

For a given *G*, we call the combination of the *c*() function and the *B*[] arrays a **graph Positional Burrows Wheeler Transform** (**gPBWT**). We submit that a gPBWT is sufficient to represent *T*, and, moreover, that it allows efficient counting of the number of threads in *T* that contain a given new thread as a subthread. Figure 2 and Table 1 give a worked example.

**Fig. 2.**
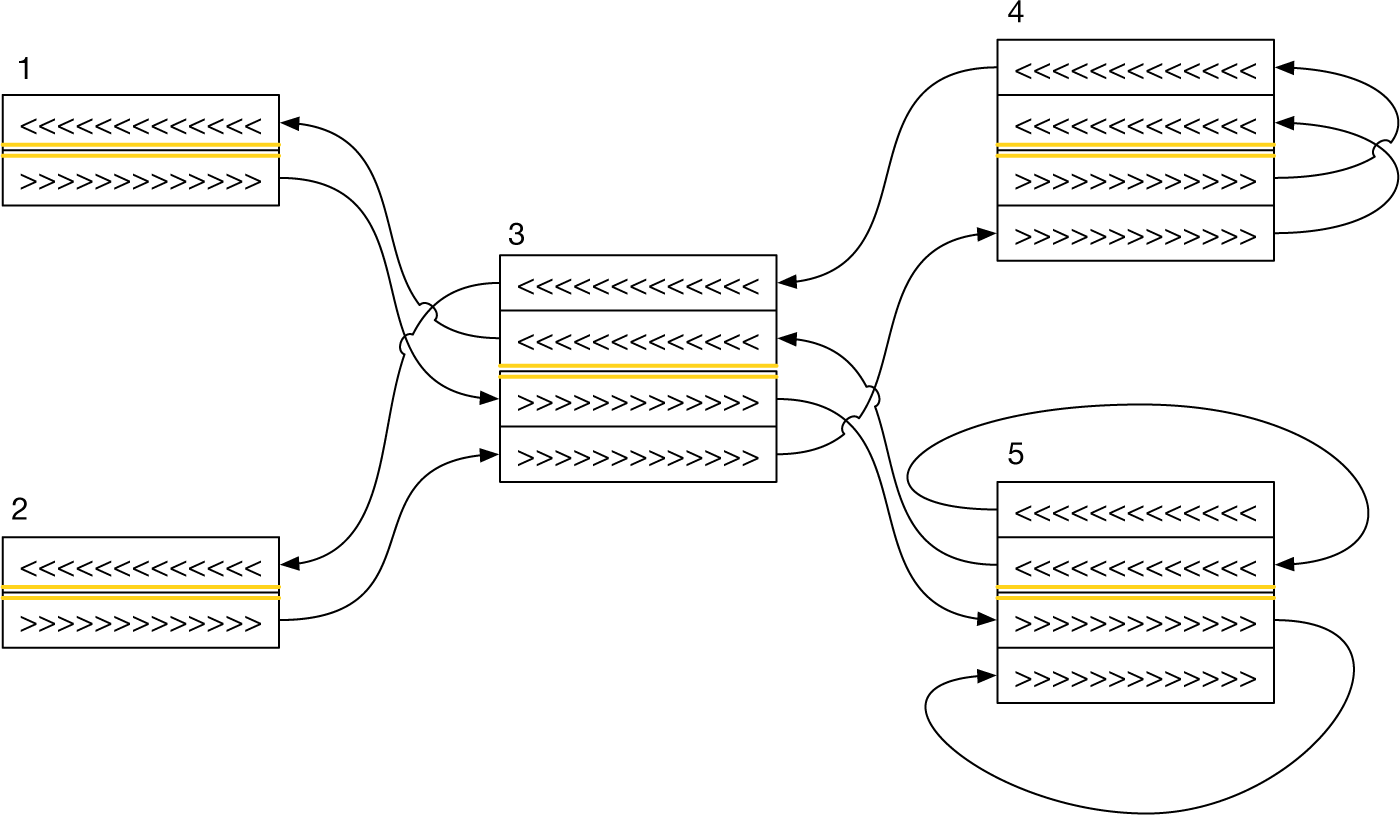
A diagram of a graph containing two embedded threads. The graph consists of nodes [1, 2, 3,4,5], with sides [1*L*, 1*R*, 2*L*, 2*R*,…], connected by edges [1*R*, 3*L*], [2*R*, 3*L*], [3*R*, 4*L*], [3*R*, 5*L*], [4*R*,4*R*], and [5*R*, 5*L*]. Embedded threads travel on the right-hand side of the nodes they are traveling through. Each thread here corresponds to a pair of “lanes” running in opposite directions. Visits are ordered from top to bottom, with “lanes” for lesser visits above those for greater ones. the “lanes” on the top half of each node are ordered in correspondence with the *B*_*s*_[] entries for the right side of the node, and those on the bottom half are ordered in correspondence with the *B*_*s*_[] entries for the right side of the node. The threads shown here are [1*L*, 1*R*, 3*L*, 3*R*, 5*L*, 5*R*, 5*L*, 5*R*] and [2*L*, 2*R*, 3*L*,3*R*, 4*L*,4*R*, 4*R*, 4*L*].

**Table 1.**
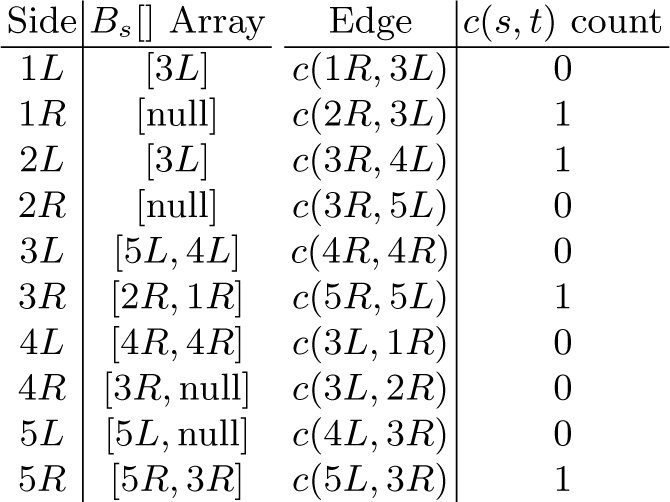
*B*_*s*_[] and *c*() values for the embedding of threads illustrated in Figure 2.

## 4 Extracting Threads

To reproduce *T* from *G*, and the gPBWT, consider each side *s* in *G* in turn. Establish how many threads begin (or, equivalently, end) at *s* by taking the minimum of *c*(*x, s*) for all sides *x* adjacent to *s*. If *s* has no incident edges, take the length of *B*_*s*_[] instead. Call this number *b*. Then, for *i* running from 0 to *b*, exclusive, begin a new thread at *n*(*s*) with the sides [*s*, 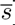]. Next, we traverse from *n*(*s*) to the next node. Consult the *B*_*s*_[*i*] entry. If it is the null side, stop traversing, yield the thread, and start again from the original node *s* with the next *i* value less than *b*. Otherwise, traverse to side *s’* = *B*_*s*_[*i*]. Calculate the arrival index *i’* as *c*(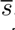, *s’*) plus the number of entries in *B*_*s*_[] before entry *i* that are also equal to *s’*. This gives the index in 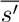 of the thread being extracted. Then append *s’* and 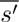 to the growing thread, and repeat the traversal process with *i* ← *i’* and *s* ← *s’*, until the end of the thread is reached.

This process will enumerate all threads in the graph, and will enumerate each such thread twice (once from each end). The threads merely need to be deduplicated (such that two enumerated threads produce one actual thread, as the original collection of embedded threads may have had duplicates) in order to produce the collection of embedded threads *T*. Pseudocode for thread extraction is shown in Algorithm 1.

### Algorithm 1 Algorithm for extracting threads from a graph.

**function** starting_AT(*Side, G, B*[], *c*())

▹ Count instances of threads starting at *Side*.

▹ Replace by an access to a partial sum data structure if appropriate.

**if** *Side* has incident edges **then**

**return** *c*(*s,Side*) for minimum *s* over all sides adjacent to *Side*.

**else**

**return** LENGTH(*B*_*Side*_[])

**function** RANK(*b*[], *Index, Item*)

▹ Count instances of *Item* before *Index* in *b*[].

▹ Replace by RANK of a rank-select data structure if appropriate.

*Rank* ⟵;0

**for** *i* ⟵ 0; *i* < *Indec i* ⟵ *i* + 1 **do**

**if** *b*[*i*] = *Item* **then**

*Rank* ⟵; *Rank* + 1

**return** *RANK*

**function** WHERE_TO(*Side, Index, B*[], *c*())

▹ For thread visiting *Side* with *Index* in the reverse prefix sort order, get the corresponding sort index of the thread for the next side in the thread.

**return** *c*(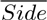, *B*_*Sid*_ [*Index*]) + Rank(*B*_*Side*_[], *Index, B*_*Side*_[*Index*])

**function** EXTRACT(*G, c*(), *B*[])

▹ Extract all oriented threads from graph *G*.

**for all** Side *s* in *G* **do**
*TotalStarting* ⟵ STARTING_AT(*s*, *G, B*[], *c*())

**for** *i* ⟵ 0; *i* < *TotalStarting*; *i* ⟵ *i* + 1 **do**

*Side* ⟵ *s*

*Index* ⟵ *i*

*Thread* ⟵ [*s*, 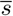]

**loop**

*NextSide ⟵ *B*_*Side*_ [*Index*]*

**if** *NextSide* = null **then**

**yield** *Thread*

**break**

*Thread* ⟵ *Thread* + [*NextSide*, 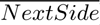]

*Index* ⟵ WHERE_TO(*Side, Index, B*[], *c*())

*Side* ⟵ *NextSide*

## 5 Succinct Storage

For the case of storing haplotype threads specifically, we can assume that, because of linkage, many threads in *T* are identical local haplotypes for long runs, diverging from each other only at relatively rare crossovers or mutations. Because of the reverse prefix sorting of the visits to each side, successive entries in the *B*[] arrays are thus quite likely to refer to locally identical haplotypes, and thus to contain the same value for the side to enter the next node on. Thus, the *B*[] arrays should benefit from run-length compression. Moreover, since (as will be seen below) one of the most common operations on the *B*[] arrays will be expected to be rank queries, a succinct representation, such as a collection of bit vectors or a dynamic wavelet tree, would be appropriate. To keep the alphabet of symbols in the *B*[] arrays small, it is possible to replace the stored sides for each *B*[] with numbers referring to the nodes adjacent to 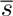

We note that, for contemporary variant collections (e.g. the 1000 Genomes Project), the underlying graph *G* may be very large, while there may be relatively few threads (of the order of thousands) [1]. Implementers should thus consider combining multiple *B*[] arrays into a single data structure to minimize overhead.

## 6 Embedding Threads

A trivial construction algorithm for the gPBWT is to independently construct *B*_*s*_[] and *c*(*s, s’*) for all sides *s* and oriented edges (*s, s’*) according to their definitions above. However, this would be very inefficient. Here we present an efficient algorithm for gPBWT construction, in which the problem of constructing the gPBWT is reduced to the problem of embedding an additional thread.

Each thread is embedded by embedding its two orientations, one after the other. To embed a thread orientation *t* = [*t*_0_, *t*_1_,…*t*_2*N*_, *t*_2*N*_+1] we first look at node *n*(*t*_0_), entering by *t*_0_. We insert a new entry for this visit into *B*_*t*0_[], lengthening the array by one. The location of the new entry is near the beginning, before all the entries for visits arriving by edges, with the exact location determined by the arbitrary order imposed on thread orientations. Thus, its addition necessitates incrementing *c*(*s,t*_0_) by one for all oriented edges (*s,t*_0_) incident on *t*_0_ from sides *s* in *G*. If no other order of thread orientations suggests itself, the order created by their addition to the graph will suffice, in which case the new entry can be placed at the beginning of *B*_*t*0_[]. We call the location of this entry *k*. The value of the entry will be *t*_2_, or, if *t* is not sufficiently long, the null side, in which case we have finished.

If we have not finished the thread, we first increment *c*(*s,t*_2_) by one for each side *s* adjacent to *t*_2_ and after *t*_1_ in the global ordering of sides. This updates the *c*() function to account for the insertion into *B*_*t*2_[] we are about to make. We then find the index at which the next visit in *t* ought to have its entry in *B*_*t*2_[], given that the entry of the current visit in *t* falls at index *k* in *B*_*t*0_[]. This is given by the same procedure used to calculate the arrival index when extracting threads, denoted as *WHERE_TO*(*t*_1_, *k*) (see Alg. 1). Setting *k* to this value, we can then repeat the preceding steps to embed *t*_2_, *t*_3_, etc. until *t* is exhausted and its embedding terminated with a null-side entry. Pseudocode for this process is shown in Algorithm 2.

Assuming that the *B*[] array information is both indexed for *O*(*log*(*n*)) rank queries and stored in such a way as to allow *O*(*log*(*n*)) insertion and update (in the length of the array *n*), this insertion algorithm is *O*(*N* · *log*(*N* + *E*)) in the length of the thread to be inserted (*N*) and the total length of existing threads (*E*). Inserting *M* threads of length *N* will take *O*(*M* · *N* · *log*(*M · N*)) time.

### Algorithm 2 Algorithm for embedding a thread in a graph.

**procedure** INSERT(*b*[], *Index, Item)*

▹ Insert *Item* at *Index* in *b*[].

▹ Replace by insert of a rank-select-insert data structure if appropriate.

LENGTH(*b*[]) ⟵ LENGTH(*b*[]) + 1 ▹ Increase length of the array by 1

**for** *i* ⟵ LENGTH(*b*[]) − 1; *i* > *Index; i* ⟵ *i* − 1 **do**

*b*[*i*] ⟵ *b*[*i* − 1]

*b*[*Index*] = *Item*

**procedure** INCREMENT_C(*Side, NextSide, c*())

▹ Modify *c*() to reflect the addition of a visit to the edge (*Side, NextSide*).

**for all** side *s* adjacent to *NextSide* in *G* **do**

if *s* > *Side* in side ordering **then**

*c*(*s, NextSide*) ⟵ *c*(*s, NextSide*) + 1

**procedure** EMBED(*t, G, B*[], *c*())

▹ Embed an oriented thread *t* in graph *G*.

▹ Call this twice to embed it for search in both directions.

*k* ⟵ 0 ▹ Index we are at in *B*_t_2i__ []

**for** *i* ⟵ 0; 2*i* < LENGTH(*t*); *i ⟵ *i* + 1 **do***

**if** 2*i* + 2 < LENGTH(*t*) **then**

▹ The thread has somewhere to go next.

INSERT(*B*_t_2i__[], *k,t*_2i_+a2)

INCREMENT_C(*t*_2i_+1, *t*_2i_+2, *c*())

*k* ⟵ WHERE_TO(*t*_2i_, *k*, *B*[], *c*())

**else**

INSERT(*B*_t_2i__ [], *k*, null)

## 7 Counting Occurrences of Subthreads

The generalized PBWT data structure presented here preserves some of the original PBWT’s efficient haplotype search properties [2]. The algorithm for counting all subthread instances in *T* of a new thread orientation *t* runs as follows.

We define ƒ_*i*_ and *g*_*i*_ as the first and past-the-last indexes for the range of visits of threads in *T* to side *t*_2i_, ordered as in *B*_t_2i__[].

For the first step of the algorithm, ƒ_0_ and *g*_*0*_ are initialized to 0 and the length of *B*_t_0__[], respectively, so that they select all visits to node *n*(*t*_0_), seen as entering through *t*_0_. On subsequent steps, ƒ_i+1_ and *g*_i+1_, are calculated from ƒ*i* and *g*_i_ merely by applying the *WHERE_TO*() function (see Alg. 1). We calculate ƒ_i+_1 = WHERE_TO(*t*_2i_, ƒ_i_) and *g*_*i*+1_ = WHERE_TO(*t*_2i_, *g*_*i*_).

This process can be repeated until either ƒ_i+1_ ≥ *g*_*i*+1_, in which case we can conclude that the threads in the graph have no matches to *t* in its entirety, or until *t*_2*N*_, the last entry in *t*, has its range ƒ*N* and *g*_*N*_ calculated, in which case *g*_N_ — ƒ*N* gives the number of occurrences of *t* as a subthread in threads in *T*. Moreover, given the final range from counting the occurrences for a thread *t*, we can count the occurrences of any longer thread that begins with *t*, merely by continuing the algorithm with the additional entries in the longer thread.

Assuming that the *B*[] arrays have been indexed for *O*(1) rank queries, the algorithm is *O*(*N*) in the length of the subthread *t* to be searched for, and has a runtime independent of the number of occurrences of *t*. Pseudocode is shown in Algorithm 3.

### Algorithm 3 Algorithm for searching for a subthread in the graph.

**function** COUNT(*t, G, B*[], *c*())

▹ Count occurrences of subthread *t* in graph *G*.

ƒ ⟵ 0

*g* ⟵ LENGTH(*B*_t_0__ [])

**for** *i* ⟵ 0; 2(*i* + 1) < LENGTH(*t*); *i* ⟵ *i* + 1 **do**

ƒ ⟵ WHERE_TO(*t*_2i_, ƒ, *B*[], *c*())

*g* ⟵ WHERE_TO(*t*_2i_, *g*, *B*[], *c*())

**if** ƒ ≥ *g* **then**

**return** 0

**return** *g* − ƒ

## 8 Results

The gPBWT was implemented within xg, the succinct graph indexing component of the vg variation graph toolkit [3]. Due to the succinct data structure libraries employed, efficient integer vector insert operations were not possible, and so a batch construction algorithm, applicable only to directed acyclic graphs, was implemented. A modified release of vg, which can be used to replicate the results shown here, is available from https://github.com/adamnovak/vg/releases/tag/gpbwt-paper.

The modified vg was used to create a variation graph for human chromosome 22, using the 1000 Genomes Phase 3 VCF on the hg19 assembly, embedding information about the correspondence between VCF variants and graph elements [1]. Next, haplotype information for the 5,008 haplotypes stored in the VCF was imported and stored in a gPBWT-enabled xg index for the graph, using the batch construction algorithm mentioned above. In cases where the VCF specified self-inconsistent haplotypes (for example, a haplotype with a G to C SNP and a G to GAT insertion at the same position), they were broken apart at the inconsistent positions. The xg indexing and gPBWT construction process took 25 hours and 45 minutes using a single indexing thread on an Intel Xeon X7560 running at 2.27 GHz, and consumed 344 GB of memory. The high memory usage was a result of the decision to retain the entire data set in memory in an uncompressed format during construction. However, the resulting xg index was 662 MB on disk, of which 573 MB was used by the gPBWT. Information on the 5,008 haplotypes across the 1,103,547 variants was thus stored in about 1.7 bits per phased diploid genotype in the succinct self-indexed representation, or 0.018 bits per haplotype base. Extrapolating linearly from the 51 megabases of chromosome 22 to the entire 3.1 gigabase human reference genome, a similar index of the entire 1000 Genomes dataset would take 40 GB, with 35 GB devoted to the gPBWT. This is well within the storage and memory capacities of modern computer systems.

### Random Walks

To evaluate query performance, 1 million random walks of 100 bp each were simulated from the graph. To remove walks covering ambiguous regions, walks that contained two or more N bases in a row were eliminated, leaving 686,897 random walks. The number of haplotypes in the gPBWT index consistent with each walk was then determined, taking 81.30 seconds in total using a single query thread on the above-mentioned Xeon system. The entire operation took a maximum of 685 MB of memory, indicating that the on-disk index did not require significant expansion during loading to be usable. Overall, the gPBWT index required 118 microseconds per count operation on the 100 bp random walks. It was found that 317,681 walks, or 46%, were not consistent with any haplotype in the graph. The distribution of of the number of haplotypes consistent with each random walk is visible in Figure 3.

**Fig. 3.**
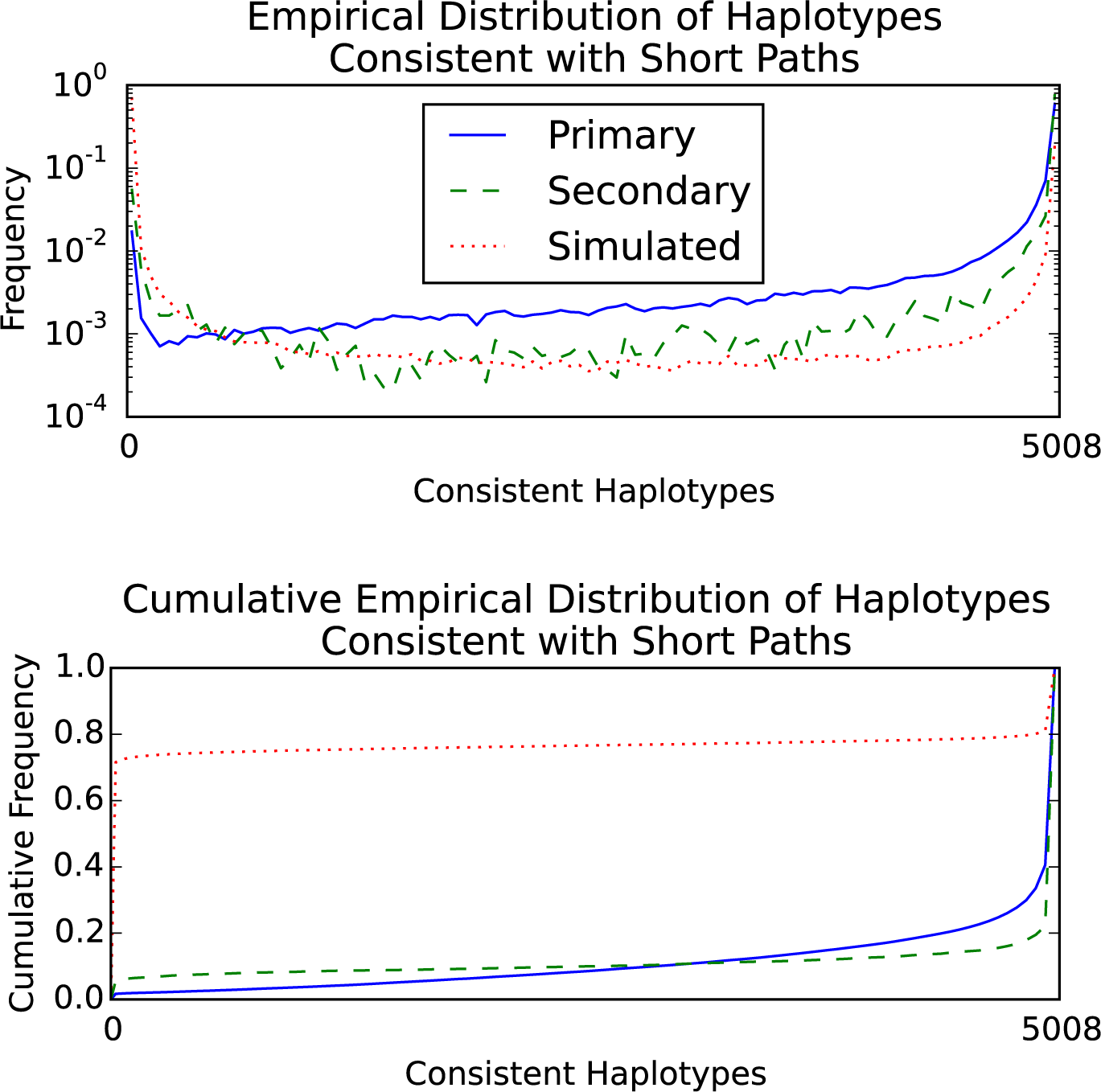
Distribution (top) and cumulative distribution (bottom) of the number of 1000 Genomes Phase 3 haplotypes consistent with short paths in the chromosome 22 graph. Primary mappings of 101 bp reads with scores of 90 out of 101 or above (*n* = 1, 509, 672) are the solid blue line. Secondary mappings meeting the same score criteria (*n* = 57, 115) are the dashed green line. Simulated 100 bp random walks in the graph without consecutive N characters (*n* = 686, 897) are the dotted red line. Consistent haplotypes were counted using the gPBWT support added to `vg` [3].

### Read Mapping

To further evaluate the performance of the query implementation, 1000 Genomes Low Coverage Phase 3 reads for NA12878 that were mapped in the official alignment to chromosome 22 were downloaded and re-mapped to the chromosome 22 graph, using the xg/GCSA2-based mapper in vg, allowing for up to a single secondary mapping per read. The reads which mapped with scores of at least 90 points out of a maximum of 101 points (for a perfectly-mapped 101 bp read) were selected (so filtering out alignnments highly like to be erroneous), and broken down into primary and secondary mappings. The number of haplotypes in the gPBWT index consistent with each read’s path through the graph was calculated (Figure 3). For 1,509,672 primary mappings, the count operation took 226.36 seconds in total, or 150 microseconds per mapping, again using 685 MB of memory. It was found that 13,918 of these primary mappings, or 0.9%, and 1,280 of 57,115 secondary mappings, or 2.2%, were not consistent with any haplotype path in the graph. These read mappings, despit having reasonable edit based scores, may represent rare recombinations, but the set is also likely to be enriched for spurious mappings.

## 9 Discussion

We have introduced the gPBWT, a graph based generalization of the PBWT. We have demonstrated that a gPBWT can be built for a substantial genome graph (all of human chromosome 22 and the associated chromosome 22 variants in 1000 Genomes), and extrapolated that a whole-genome gPBWT could be constructed for all 5,008 haplotypes of the 1000 Genomes data and stored in the main memory of a contemporary computer. Looking forward, this combination of genome graph and gPBWT could potentially enable efficient mapping not just to one reference genome or collapsed genome graph, but simultaneously to a large set of genomes related by a genome graph.

## 10 Acknowledgements

We would like to thank Richard Durbin for inspiration and David Haussler for his extremely helpful comments on the manuscript.

